# Crosstalk between age accumulated DNA-damage and the SIRT1-AKT-GSK3ß axis in urine derived renal progenitor cells

**DOI:** 10.1101/2022.05.07.491023

**Authors:** Lars Erichsen, James Adjaye

## Abstract

The aging process is manifested by a multitude of interlinked biological processes. These processes contribute to genomic instability, telomere attrition, epigenetic alterations, loss of proteostasis, deregulated nutrient-sensing, mitochondrial dysfunction, cellular senescence, stem cell exhaustion, and altered intercellular communication. Together these are recognized as of the main risk factors of the world’s most prevalent diseases, such as neurodegenerative disorders, cancer, cardiovascular disease, and metabolic disease.

The mammalian ortholog of the yeast silent information regulator (Sir2) SIRT1 is a NAD^+^-dependent class III histone deacetylase and has been recognized to be involved in many of the forementioned processes. Therefore, its activity is connected to aging via the regulation of apoptosis, cell differentiation, development, stress response, metabolism, and tumorigenesis. Furthermore, the physiological activity of several sirtuin family members has been connected to the regulation of life span of lower organisms (Caenorhabditis elegans and Drosophila melanogaster) as well as mammals. Aging in somatic cells of mammals is accompanied by mutations and other forms of DNA damage. These might manifest in transient cell cycle arrest associated with DNA repair, apoptosis, senescence, or cell differentiation. The activity of SIRT1 has previously been reported to be regulated by the DNA damage response pathway. On the one hand, SIRT1 is recruited from ATM to DBS and is required for DNA damage repair, but on the other hand, SIRT1 activity was also found to be negatively regulated by genotoxic stress via the interaction of ATM with Deleted in Breast Cancer 1 (DBC1). Increased levels of DBS are associated with downregulation of ATM and lower phosphorylation levels of AKT and GSK3ß, with significant implications for mesenchymal stem cell (MSC) maintenance and differentiation. In this proposed “stem cell checkpoint,” the ATM signalling pathway initiated by DBS maintains MSCs and blocks their differentiation. Based on this, it has already been established that in senescent mesenchymal stem cells, SIRT1 expression is decreased, while its overexpression delays the onset of senescence and loss of differentiation capacity/ability.

In the present study, we provide evidence that SIX2-positive urine derived renal progenitor cells-UdRPCs isolated directly from human urine show typical hallmarks of aging when obtained from elderly donors. This includes the transcriptional downregulation of SIRT1 and its downstream targets AKT and GSK3ß. This transcriptional downregulation is accompanied by an increase in DNA damage and transcriptional levels several cell cycle inhibitors such as P16, reflecting possibly the ATM induced “stemness checkpoint” to maintain UdRPC stemness and differentiation capacity. We provide evidence that the renal progenitor transcription factor SIX2 binds to the coding sequence of SIRT1 and both factors mutually influence the transcription of each other. Furthermore, we show that the SIRT1 promoter region is methylation sensitive and becomes subsequently methylated in UdRPCs derived from aged donors, dividing them into SIRT1 high and low expressing UdRPCs. This downregulation might render the cells more vulnerable to endogenous noxae accelerating the accumulation of DNA damage and ultimately the accumulation of aging associated hallmarks.

## Introduction

Recent demographic studies suggest a considerable increase in the number of elderly people within the next decades [2]. The aging process has been recognized as one of the main risk factors of the world’s most prevalent diseases, including neurodegenerative disorders, cancer, cardiovascular disease and metabolic disease [3]. Aged tissues are characterized by a progressive loss of physiological integrity, leading to impaired function and increased vulnerability to death. On molecular level, Otin et al., proposed nine candidate hallmarks of aging, which are considered to contribute to the aging process in general and collectively contribute to the aging phenotype [4]. In detail these hallmarks are: genomic instability, telomere attrition, epigenetic alterations, loss of proteostasis, deregulated nutrient-sensing, mitochondrial dysfunction, cellular senescence, stem cell exhaustion, and altered intercellular communication [4].

As Mammals age their cells accumulate somatic mutations and other forms of DNA damage, such as chromosomal abnormalities and changes in chromosome copy number [5]. When these alterations arise the cell cycle is arrested in G1 phase, mainly triggered by the activity of TP53 and/or TP16 [6–8]. Depending on the cell type, an active DNA damage response (DDR) has four potential outcomes, namely, transient cell cycle arrest coupled with DNA repair, apoptosis, senescence or cell differentiation [9]. For example, it is well established that ataxia telangiectasia mutated (ATM) [10] and TP53 [11, 12] are capable of regulating hemopoietic stem cell quiescence or senescence and self-renewal. Furthermore, both show a biphasic response to DNA-damage in a dose dependent manner [12, 13]. A major pathway that becomes activated by the DDR is the phosphatidylinositol-3-kinase/protein kinase B pathway (PI3K/AKT) [14]. It has been recognized that ATM and DNA-dependent protein kinase (DNA-PKs) are involved in AKT activation at the site of double strand breaks and inhibition of AKT activity impairs the repair of DNA double-strand breaks (DBS) [15–17]. On the other hand, AKT is also activated by growth factors and promotes cell cycle progression at G1/S and G2/M transition [14, 18, 19].

SIRT1 is the mammalian ortholog of the yeast silent information regulator (Sir2) and as a NAD^+^-dependent class III histone deacetylase involved in many processes connected to aging, like apoptosis, cell differentiation, development, stress response, metabolism, and tumorigenesis [20–22]. The high number of cellular features that can be regulated by SIRT1 is based on its variety of target molecules. Beside its specificity for the histone proteins H1, H3 and H4 and thereby promoting the formation of heterochromatin and transcriptional repression, SIRT1 has been reported to also deacetylate several transcription factors [23–25], and apoptosis and cell-cycle regulating proteins, including TP53 [26, 27]. The physiological activity of several sirtuin family members has been connected to the regulation of life span of lower organisms such as Caenorhabditis elegans [1], and Drosophila melanogaster [28] as well as mammals [29]. While SIRT1 is recruited to DBS by ATM and is required for DNA damage repair [30], it has also been noticed that SIRT1 activity is negatively regulated by genotoxic stress via ATM interaction with deleted in breast cancer 1 (DBC1) [31, 32]. In senescent mesenchymal stem cells (MSCs) SIRT1 expression is reduced, while its over-expression delays the onset of senescence and the loss of differentiation capacity [33].

We recently reported human urine as a non-invasive source of renal stem cells with regenerative potential [34]. The urine derived renal progenitor cells (UdRPCs) express renal stem cell markers such as SIX2, CITED1 WT1, CD133, CD24 and CD106. Here we provide evidence that SIX2-positive urine derived renal progenitor cells-UdRPCs isolated from human urine show typical hallmarks of aging when obtained from elderly donors. This includes the transcriptional downregulation of SIRT1 and its downstream targets AKT and GSK3ß. This transcriptional downregulation is accompanied by an increase in DNA damage and transcriptional levels several cell cycle inhibitors such as P16. We provide evidence that the renal progenitor transcription factor SIX2 binds to the coding sequence of SIRT1 and both factors mutually influence the transcription of each other. Furthermore, we show that the SIRT1 promoter region is methylation sensitive and becomes subsequently methylated in UdRPCs derived from aged donors, dividing them into SIRT1 high and low expressing UdRPCs. We propose the SIRT1-AKT-GSK3ß axis to regulate and monitor self-renewal capacity of urine derived renal progenitor cells.

## Material and Methods

### Cell culture conditions

UdRPCs were cultured in Proliferation Medium (PM) composed of 50% DMEM high Glucose and 50% Keratinocyte medium supplemented with 5% FCS, 0.5% NEAA, 0.25% Gtx and 0.5% Penicillin and Streptomycin at 37°C (Gibco, Carlsbad, United States) under hypoxic conditions. For all experiments cells were collected after 7-8 passages and seeded in 6-or 12-well plates coated with 0.2% Gelatin (Thermo Fisher Scientific, Waltham, United States). Resveratrol (Sigma Aldrich, St. Louis, Missouri, United States) and Bleomycin (Sigma Aldrich) were added to the the culture medium to a final concentration of 30 μg/ml. Cells were incubated with Resveratrol and Bleomycin containing culture medium for 24h.

### Relative Quantification of aging-associated gene expression by real-time PCR

Total RNA was extracted from UdRPCs using the RNeasy Mini Kit (Qiagen, Hilden, Germany) according to the manufacturer’s instructions. First-strand cDNA synthesis was performed from 1 μg RNA by reverse transcription using oligo(dT) (Promega, Madison, United States) and Moloney murine leukemia virus reverse transcriptase (Promega) in a volume of 50 μL at 42 °C for 1 h.

Real time PCR of aging associated gene expression was performed as follows:

Real time measurements were carried out on the Step One Plus Real Time PCR Systems using MicroAmp Fast optical 384 Well Reaction Plate and Power Sybr Green PCR Master Mix (Applied Biosystems, Foster City, United States). The amplification conditions were denaturation at 95°C for 13 min. followed by 37 cycles of 95°C for 50s, 60°C for 45s and 72°C for 30s. Primer sequences are listed in supplement table 1.

### Immunofluorescence staining

Cells were fixed with 4% paraformaldehyde (PFA) (Polysciences, Warrington, United States). Unspecific binding sites of the fixed cells were blocked by incubation with blocking buffer containing 10% normal goat or donkey serum, 1% BSA, 0.5% Triton, and 0.05% Tween, for 2h at room temperature. The primary antibody was diluted 1:1 in blocking buffer with PBS and incubated at 4°C overnight (or at least 16h). After incubation the cells were washed three times with PBS/0.05% Tween and the secondary antibodies were diluted the same way as the primary antibodies with a 1:500 dilution. After 1h of secondary antibody incubation the cells were washed again three times with PBS/0.05% Tween and nuclei were stained with Hoechst 1:5000 (Thermo Fisher Scientific) and cytoskeleton was stained with Alexa Flour 488 phalloidin (Thermo Fisher Scientific) (1:400). Images were captured using a fluorescence microscope (LSM700; Zeiss, Oberkochen, Germany) with Zenblue software (Zeiss). Individual channel images were processed with Fiji. Detailed Information of the used antibodies are given in supplementary table 2.

### Microarray data analyses

Total RNA (1 μg) preparations were hybridized on the PrimeView Human Gene Expression Array (Affymetrix, Thermo Fisher Scientific, United States) at the core facility Biomedizinisches Forschungszentrum (BMFZ) of the Heinrich Heine University Düsseldorf. The raw data was imported into the R/Bioconductor environment and further processed with the package affy using background-correction, logarithmic (base 2) transformation and normalization with the Robust Multi-array Average (RMA) method. The heatmap.2 function from the gplots package was applied for cluster analysis and to generate heatmaps using Pearson correlation as similarity measure. Gene expression was detected using a detection-p-value threshold of 0.05. Differential gene expression was determined via the p-value from the limma package which was adjusted for false discovery rate using the q value package. Thresholds of 1.33 and 0.75 were used for up-/down-regulation of ratios and 0.05 for p-values. Venn diagrams were generated with the Venn Diagram package. Subsets from the venn diagrams were used for follow-up GO and pathway analyses as described by Zhou et al [35]. Gene expression data will be available online at the National Centre of Biotechnology Information (NCBI) Gene Expression Omnibus.

### Western blot analysis

UdRPCs were lysed in lysis buffer composed of 5 M NaCl, 1% NP-40, 0.5% DOC, 0.1% SDS, 1 mM EDTA, 50 mM Tris, pH 8.0, and freshly added 10 μL/mL protease- and phosphatase inhibitor (Sigma Aldrich). 20 μg of the obtained protein lysate was resolved in a 10% sodium dodecyl sulfate-PAGE gel and transferred onto Immobilon-P membrane (Merck Millipore, Burlington, United States). Membranes were probed with primary antibody at 4 °C overnight, washed three times with 0.1% Tween-20 in Tris-buffered saline, and incubated with secondary antibody for 1h at room temperature. The signals were visualized with enhanced luminescence Western Bright Quantum (Advansta, Bering Dr, United States). Detailed Information of the used antibodies are given in supplementary table 2.

### Expression of Lamin A and Progerin

Total RNA was extracted from UdRPCs using the RNeasy Mini Kit (Qiagen, Hilden, Germany) according to the manufacturer’s instructions. First-strand cDNA synthesis was performed from 1 μg RNA by reverse transcription using oligo(dT) (Promega, Madison, United States) and Moloney murine leukemia virus reverse transcriptase (Promega) in a volume of 50 μL at 42 °C for 1 h. Lamin A and Progerin were detected as described by <u>McClintock</u> et al., [36]. Primer sequences are listed in supplement table 1.

### Immunoprecipitation

Cells were chemically crosslinked with 11% formaldehyde solution for 15 min at room temperature. Cells were washed twice with 1× PBS and harvested using a silicon scraper in a lysis buffer, and genomic DNA was sonicated at 4 °C in TPX® polymethylpentene tubes using a Bioruptor® sonicator (Diagenode, Liege, Belgium). Twenty sonication pulses of each 15 sec were applie. The resulting wholecell extract (WCE) was incubated overnight at 4°C with 100 µl of Dynal Protein A magnetic beads (Diagenode) previously pre-incubated with (input) and without (negative control) 10 µg of SIX2 antibody. Beads were washed five times with RIPA buffer and once with TE containing 50 mM NaCl. Bound complexes were eluted from the beads by heating at 65°C with occasional vortexing, and crosslinking was reversed by overnight incubation at 65°C. Input and negative control were also treated for crosslink reversal. Immunoprecipitated DNA and whole-cell extract DNA were then purified by treatment with RNase A, proteinase K, multiple phenol:chloroform:isoamyl alcohol extractions and precipitation with ethanol. Purified DNA was amplified using the PCR protocol.

### Bisulfite sequencing

Bisulfite sequencing was performed following bisulfite conversion with the EpiTec Kit (Qiagen, Hilden, Germany) as described in Erichsen et al. [37]. PCR primer sequences are given in supplementary table 1 and refer to +1 transcription start of the following sequences:

Homo sapiens sirtuin 1 (SIRT1), RefSeqGene on chromosome 10

NCBI Reference Sequence: NG_050664.1

Obtained sequences were analysed using Quma (http://quma.cdb.riken.jp/) as described in [38].

### Statistical analysis

Data is presented as arithmetic means + standard error of mean. At least three experiments were used for the calculation of mean values. To address the statistical significance, we applied the two-samples Student’s t-test with a significance threshold 0.05. The level of significance was set to p < 0.05.

## Results

### UdRPCs from aged donors show typical hallmarks of aging

We recently reported MSCs isolated directly from urine samples. These cells express the renal stem cell markers SIX2, CITED1 WT1, CD133, CD24 and CD106, we referred to these cells as urine derived renal progenitor cells (UdRPCs) [34]. In this study, the progenitor cells were isolated from distinct individuals of mixed ethnicity with ages ranging from 21 to 77 years. It is well documented in the literature, that MSCs show decline of self-renewal capacity and of immunosuppressive properties with increased donor age and in vitro expansion [39–42]. As previously reported UdRPCs can be kept in culture for up to 12 passages, whereas cells from aged donors show a decline of proliferation capacity after 9-10 passages. Therefore, all experiments were carried out with UdRPCs after 7-8 passages. Hierarchical clustering analysis comparing the transcriptomes of UdRPCs revealed a distinct expression pattern of cells derived from donors aged between 21 to 51 years (young, green box) and 54 to 61 years (aged, red box) (fig. 1a). Microarray analysis revealed a common expressed gene-set of 11917 between UdRPCs derived from young- and elderly donors, while 750 genes were exclusively expressed in cells derived from young- and 155 genes in cells derived from elderly individuals by comparing the expressed gene set (det-p < 0.05) (fig. 1b). The most over-represented GO BP-terms common expressed in UdRPCs derived from young- and elderly donors are associated with metabolic processes such as, organic acid transport and regulation of ion transport as well as cell junction organization and cell morphogenesis involved in differentiation. The most over-represented GO BP-terms exclusive to the young UdRPCs are associated with DNA-replication, mesodermal cell differentiation, renal system development and PI3K-AKT signalling pathway. In comparison the most over-represented GO BP-terms exclusive to aged UdRPCs are associated with assembly of collagen fibrils and other multimeric structures and regulation of calcium ion transport (fig. 1c). Additionally, we analysed our data-set for genes associated with the hallmarks of aging as proposed by Otin et al., [4]. The heatmap reveals a negative correlation between the donor age and the expression of genes involved in genomic instability (*ATM, MSH6, TFEB* and *XRCC1*), epigenetic alterations (*SETDB1, KDM6A, EZH1, SIRT1, SETDB2, HDAC6, HDAC4, KDM4B* and *SIRT3*) and genes involved in de-regulation of nutrient sensing of the one carbon-, cysteine- and methionine-metabolic pathways (*SHMT1, SHMT2* and *MAT2B*). Furthermore, our transcriptome data reveals a positive correlation between donor age and the expression of genes involved in cellular senescence (*CDKN1A, CDKN2A* and *CDKN2D*) and stem cell exhaustion (*CXCL1, IL6* and *IL8*).

**Figure 1:**
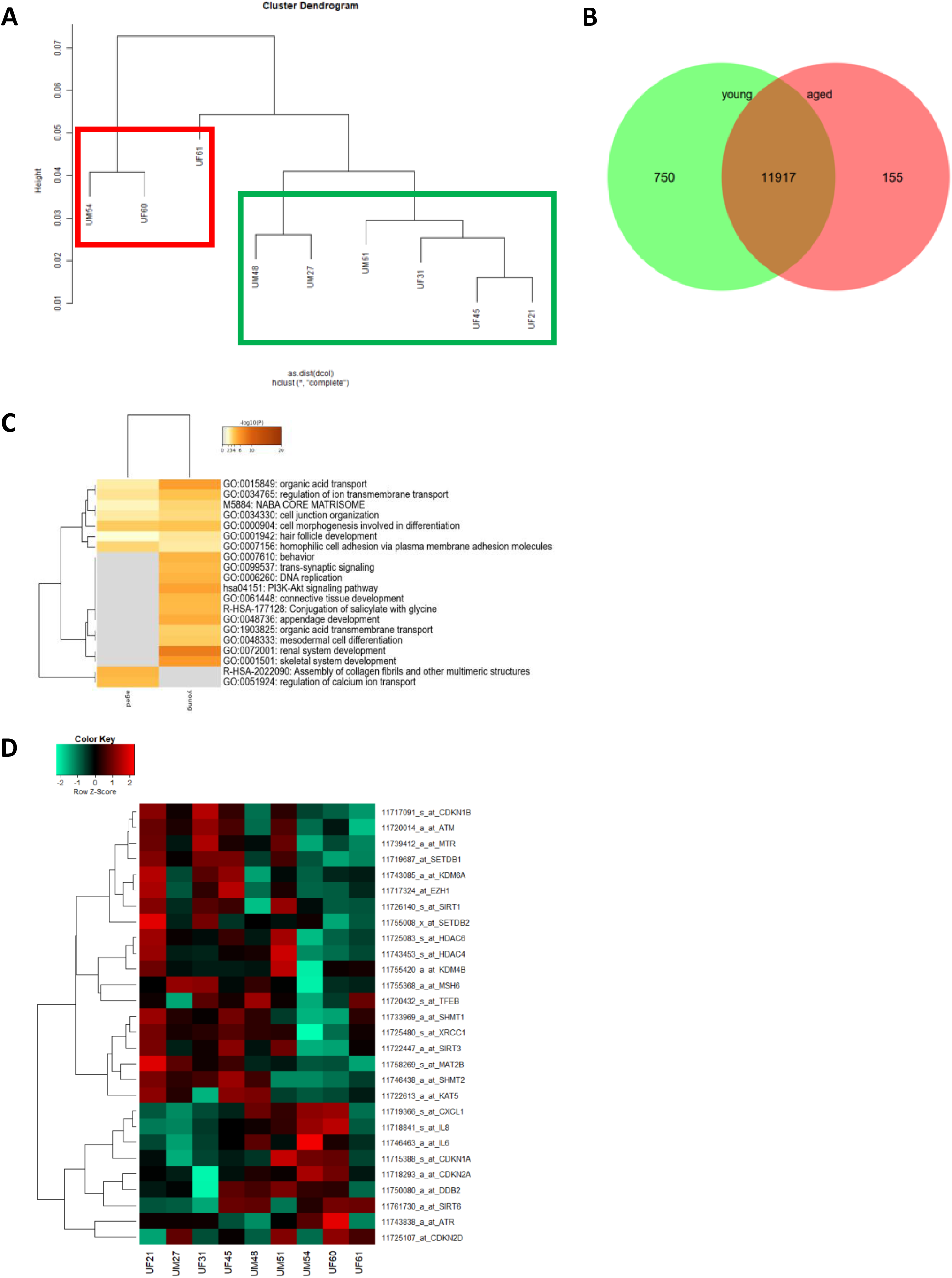
Comparative transcriptome and Gene Ontology analysis of urine-derived renal progenitors from young and aged donors. A hierarchical cluster dendrogram revealed distinct clusters of UdRPCs derived from young and aged donors (A). Expressed genes (det-p < 0.05) in UdRPCs and podocytes compared in the Venn diagram (B), shows distinct (750 in young and 155 in aged UdRPCs) and overlapping (11917) gene expression patterns. The most over-represented GO BP-terms exclusive in either young or old UdRPCs are shown in C and including DNA replication and renal development for the young UdRPCs and assembly of collagen fibrils and other multimeric structures and regulation of calcium ion transport for the old UdRPCs. The heatmap (D) reveals a distinct expression pattern between young and old UdRPCs for genes encoding for the following hallmarks of aging: genomic instability, epigenetic alterations, deregulation of nutrient sensing, cellular senescence, and stem cell exhaustion.

### UdRPCs show decline of stem cell characteristics and an increase inf DNA-damage with increase donor age

Since our microarray data revealed genes encoding for mesodermal cell differentiation and renal system development to be exclusively expressed in UdRPCs derived from young donors, we assumed that this is caused by an age-associated decline of self-renewal capacity. To test this hypothesis, we applied immunofluorescent staining for the renal stem cell marker SIX2 and qRT-PCR analysis for the stem cell markers *SIX2* and *CD133*. Surprisingly, the immunofluorescent staining revealed that almost 100% of the isolated cells from donors aged between 21 and 51 years are positive for SIX2 (fig. 2b). In contrast, qRT-PCR analysis revealed a significant downregulation/ 0.98-fold (p < 0.01) of *SIX2* mRNA expression between cells derived from the 21-year-old donor compared to cells derived from all other donors. Excluding the 21-year-old sample from the analysis, also revealed a significant downregulation/ 0.8-fold (p ≤ 0.01) of *SIX2* mRNA expression between UdRPCs derived the 27-year-old donor to all other UdRPCs. For the stem cell marker *CD133* qRT-PCR analysis revealed a significant downregulation/ 1.16-fold (p ≤ 0.01) of mRNA expression between cells derived from donors aged between 21 and 48 compared to individuals aged between 51 and 77 years (fig. 2b). As a further marker of premature terminal differentiation and/or senescence [43] we assessed the truncated form of the Lamin A transcript Progerin by semi-quantiative PCR. Strikingly, we found an increase of truncated Lamin A transcript Progerin within UdRPCs derived donors aged between 21 and 48 compared to individuals aged between 51 and 77 years) (fig. 2c). Accumulation of Progerin has been described to lead to DNA-damage and chromosomal aberrations [44, 45], by inhibiting *inter alia* the SIRT6 mediated DNA-damage repair mechanism [46]. To test our hypothesis that the identified increase of progerin mRNA in the aged UdRPCs is accompanied by increased DNA-damage, we analysed phosphorylation levels of Histone 2A (pH2A.X), an established biomarker of DNA-damage at DBS [47], by immunofluorescent based detection (fig. 2d). UdRPCs derived from individuals aged between 21 and 45 years showed a positive pH2A.X staining only in a small percentage of cells, 2% and 6% respectively. In contrast we detected a significant (p = 0.03) increase of DBS in cells derived from donors aged 51, 63 and 77 years, with 28%, 50% and 55% of cells being positive for the pH2A.X staining (fig. 2d and supplementary fig. 1a). Furthermore, we applied qRT-PCR analysis for *ATM, P16* and *TP53* (fig. 2e and supplementary fig. 1b). mRNA expression of *CDKN2A* and *TP53* were found to be not significantly altered between UdRPCs derived the different donors (p-value P16: p = 0.91 and TP53: p = 0.41), with a trend for *P16* being up- and *TP53* being downregulated with increased donor age but could also reflect heterogeneity in-between individuals. In contrast, qRT-PCR analysis revealed a significant downregulation/ 0.85-fold (p < 0.05) of *ATM* mRNA expression between cells derived from the 21-years-old donor compared to UdRPCs derived from individuals aged between 27 and 77 years. Finally, we evaluated the expression of Methionine Adenosyltransferase 2B (*MAT2b*) by qRT-PCR. This enzyme catalysizes the final step of one carbon metabolism by forming S-Adenosylmethionine from methionine and adenosine triphosphate. For *MAT2b* a significant downregulation/ 1.15-fold of mRNA levels were found in the UdRPCS derived from donors aged between 21 and 63 compared to donors aged 69 to 77 (p < 0.01) (supplementary figure. 1b).

**Figure 2:**
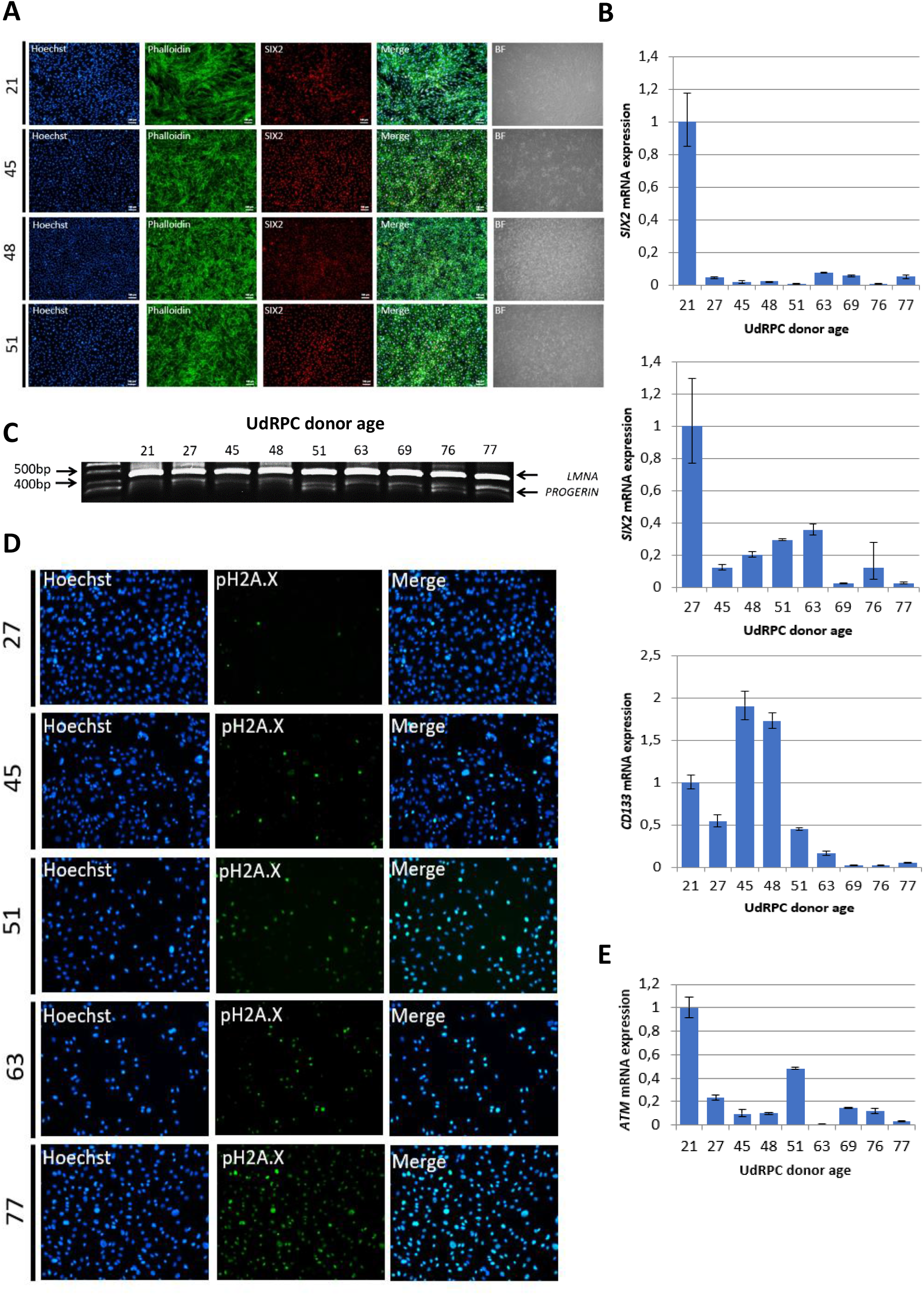
UdRPCs show decline of mesenchymal stem cell characteristics and an increase of DNA-damage with increase donor age. UdRPCs from donors aged between 21 and 77 years were isolated and stemness status was confirmed by immunofluorescent staining for the renal progenitor marker SIX2 (A) (scale bars: 100 µm). mRNA expression of SIX2 and the stem cell proliferation marker *CD133* was determined by quantitative real time PCR (B). RT-PCR analysis reveal progerin transcripts in UdRPCs. UdRPCs from donors aged between 21 and 77 years were isolated and DNA damage was visualized by immunofluorescence-basedt staining for phosphorylated Histone2A.X (pH2A.X) (D). mRNA expression of *ATM* was determined by quantitative real time PCR (E).

### Age associated changes in the SIRT1-AKT-GSK3b regulatory axis

SIRT1 [30] and the AKT pathway [14] are involved in DNA-damage-repair and our results indicate a downregulation of *SIRT1* and members of the AKT pathway in UdRPCs derived from individuals aged 48 years and above. Therefore, we evaluated the expression of SIRT1 as well as the phosphorylation levels of H2A.X, AKT and its downstream target GSK3b by Western blot detection (fig. 3a). Consistently, we found SIRT1 expression exclusively in the UdRPCs derived from individuals aged ≤ 48 years. Normalized to ß-actin expression we found a significant reduction of SIRT1 protein expression (p = 0.01) in UdRPCs derived from elderly donors by 81.3% and 64.7% (48 and 51 years). Our western blot results reveal a similar observation for the phosphorylation levels of AKT and GSK3b. Normalized to ß-actin expression we found a significant reduction of AKT (p = 0.05) and GSK3b (p = 0.04) protein phosphorylation in UdRPCs derived from donors aged 48 and 51 years, by 97.2% and 97.1% for AKT and 36.9% und 37% for GSK3b. Furthermore, we also evaluated H2A.X phosphorylation and observed an increase of DBS only in the sample derived from the 51-year-old individual. Normalized to ß-actin the increase was found to be not significant (p = 0.47) but became highly significant when H2A.X phosphorylation was normalized to the detected SIRT1 protein expression (p ≤ 0.01). To further confirm that SIRT1 becomes downregulated in UdRPCs derived from aged donors, we applied qRT-PCR analysis. According to the genecards database the *SIRT1* mRNA has six major splicing variants. Almost all splicing variants consist of exon 4 to exon 6 and can be distinguished into two groups by the existence of exon 1-3 or exon 7-10 (supplementary fig. 2). To evaluate which is the major splicing variant that changes in UdRPCs obtained from elderly donors, we designed primers that anneal in exon 1 and 2 as well as exon 7 and 8 (supplementary table 1). Surprisingly, we found both variants to be expressed and significantly altered (Exon 1-2: p = 0.03 and Exon 7-8: p = 0.01) between cells derived from the 21- and the 27-year-old donor compared to the UdRPCs derived from donors aged between 45 and 77 years, with fold changes of 0.54 and 0.75 respectively (fig. 3b). It has been recognized, that during the aging process “de novo” DNA methylation occurs within the promoter region of transcriptional downregulated genes [48, 49]. By applying genomic bisulfite sequencing we analysed the methylation status of the SIRT1 gene in UdRPCs derived from the 27-year- and the 51-year-old individual. In total a 341bp fragment of the SIRT1 promoter containing 45 CpG-dinucleotides were analysed. Stinkingly, we found 15.2% of CpGs to be methylated in the UdRPCs derived from the 27-year-old individual, while UdRPCs derived from the 51-year-old individual showed 61.3% of CpGs to be methylated. We concluded that UdRPCs could be distinguished in high and low SIRT1 expressing cells, depending on the donor age with a threshold of 48 years from our cohort. Finally, we wanted to investigate whether the age associated downregulation of SIX2, which is needed to maintain renal progenitor cells during kidney organogenesis [34], affects the level of SIRT1 mRNA expression rate. Since the UdRPCs derived from the 27-year-old individual showed the highest SIRT1 protein expression in Western blot analysis, we choose this cell line for Immunoprecipitation analysis. SIX2 has been reported to regulate the expression of Odd-skipped related 1 (Osr1) [50] and Glial Cell Derived Neurotrophic Factor (GDNF) [51] so we chose these as positive controls, while RPL0 was chosen as negative control. PCR analysis was performed with DNA derived from the whole cell extract (Input), after Immunoprecipitation with (IP) and without (negative control) antibodies against SIX2. Hereby, we could confirm a direct interaction between the SIX2 transcription factor and the genomic DNA of GDNF, OSR1 and SIRT1 (fig. 3d).

**Figure 3:**
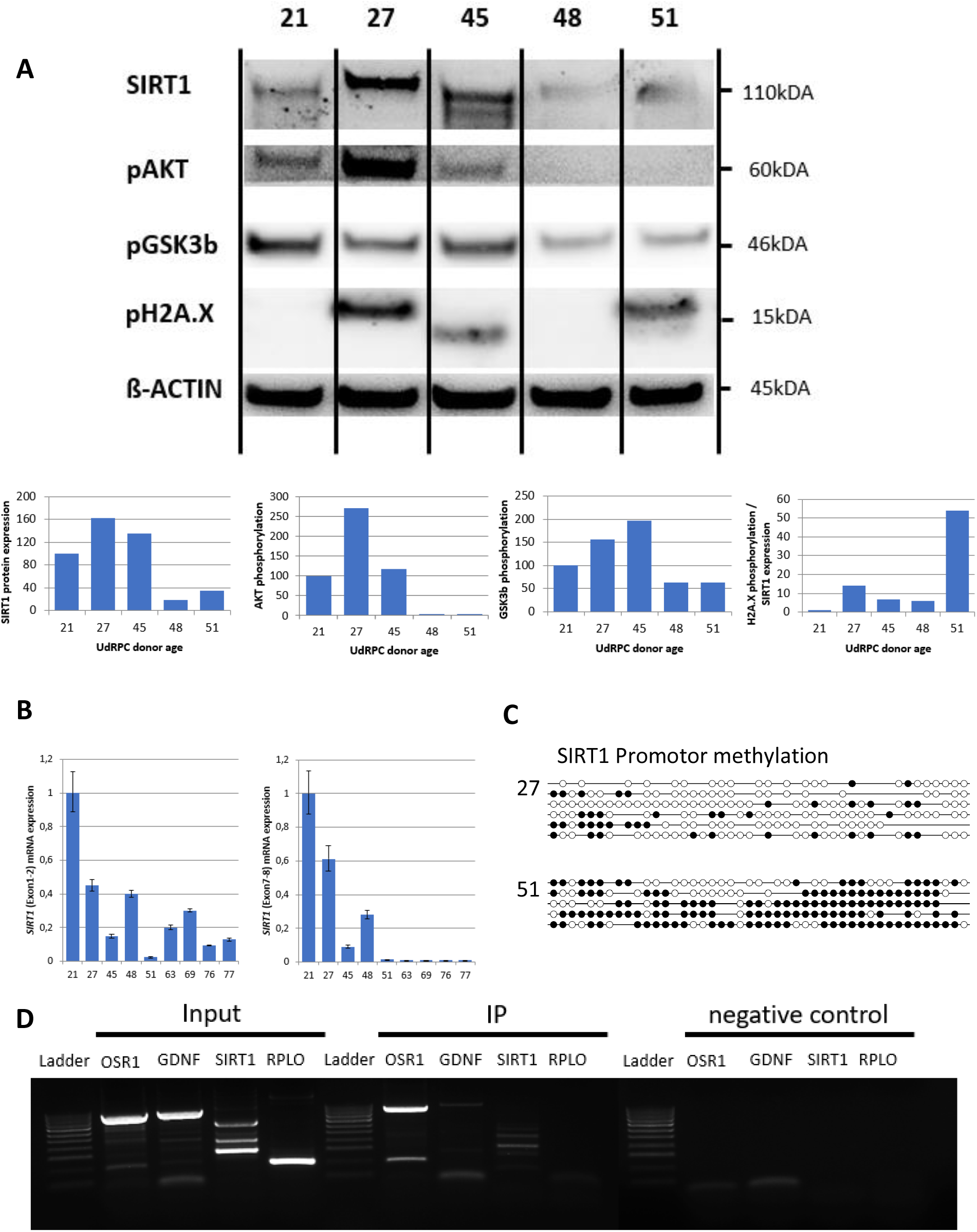
SIX2/SIRT1/AKT/GSK3b network is altered in UdRPCs derived from aged donors. Relative protein expression normalized to ß-ACTIN for SIRT1 and relative protein phosphorylation for AKT, GSK3b and pH2A.X was detected by Western blot (A). mRNA expression of *SIRT1* was determined by quantitative real time PCR (B). Detailed analyses by bisulfite sequencing of CpG island methylation patterns within 5′ regulatory region of *SIRT1* gene in young and aged UdRPCs (C). SIX2 binding within the SIRT1 gene was confirmed by Immunprecipitation followed by PCR analysis (D).

### DNA damage dependent activation of the SIX2/SIRT1/AKT/GSK3b network by resveratrol

Resveratrol has been recognized as a potent activator of SIRT1 [52, 53]. Therefore, we evaluated by qRT-PCR the effect of varying concentrations of resveratrol on low SIRT1 expressing UdRPCs with and without age-associated DNA damage. We prepared final concentrations of resveratrol directly in the cell culture medium ranging from 0.1μM to 250μM and treated the UdRPCs derived from the 48- and 51-year-old individual for 24h. Strikingly we detected a significant increase of *SIRT1* Exon 7-8 mRNA (48: p = 0.05; 9-fold and 51: p = 0.04; 0.44-fold) and SIRT1 Exon 1-2 only in the UdRPCs derived from the 51-year-old-individual (p < 0.01; 0.4-fold) (fig. 4b+f and supplementary fig. 3a). In contrast, the resveratrol treatment caused a significant downregulation of *SIRT1* Exon 1-2 mRNA in the UdRPCs derived from the 48-year-old-individual (p = 0.02; 0.52-fold) (supplementary fig. 3a). Immunofluorescence-based detection and Western blot analysis revealed a significant upregulation of SIRT1 protein by 50% and 120% respectively (48: p = 0,04 and 51: p = 0,03) for both cell lines treated with 1μM resveratrol (fig. 3a+d+h + supplementary fig. 3a). In contrast a significant downregulation of *SIRT1* mRNA (48: p < 0,01 and 51: p < 0,01) and protein (48: p < 0,01) was detected within the cells treated with the resveratrol solutions containing 2,5μM and higher concentrations by 100 and 50% respectively (fig. 4d+h and supplementary fig. 3a). To test if this effect is due to the resveratrol, we also measured the expression level of *MAT2B*. The promoter region of this gene has been identified to harbors two resveratrol binding pockets and gets activated by resveratrol in a time- and dose-dependent manner [54]. As expected *MAT2B* mRNA expression levels were significantly upregulated (48: p < 0,01 / 1.2-fold change and 51: p < 0,01 / 1.24-fold change) in the UdRPCs treated with low-concentrations and significantly downregulated (48: p < 0,01 / 0.97-fold change and 51: p < 0,01 / 0,.27-fold change) in the cells treated with the high concentrations of resveratrol. These results correlate with the observed changes in *SIRT1* mRNA expression (supplementary fig. 3b). Furthermore, immunofluorescence-based and Western blot detection of pH2A.X revealed no DBS in UdRPCs derived from the 48-year-old individual under control conditions and in the cells treated with the 1μM resveratrol solution, but a significant increase of DBS (p < 0,01; 86% of cells were found to be pH2A.X positive) was detected when cells were treated with the 2,5μM resveratrol solution (fig. 4a+d). Accordingly, 54% of UdRPCs obtained from the 51-year-old individual revealed a positive pH2A.X staining under control conditions, which became significantly elevated (51: p = 0,05; 71% of cells were found to be pH2A.X positive) by the 2,5μM resveratrol solution (fig. 4a+h). Strikingly only 34% of cells treated with 1μM resveratrol were positive for pH2A.X expression (51: p = 0,05). To test our hypothesis if resveratrol induced activation of *SIRT1* prevents cellular senescence by increasing DNA damage repair mechanisms, we evaluated mRNA expression levels of *ATM* and the cell cycle regulator *P16* by qRT-PCR (fig. 4b+f+supplementary fig. 3b). We found a significant upregulation / 26-fold of mRNA expression in the UdRPCs derived from the 48-year-old donor treated with low concentrations and a significant downregulation/ 0.99-fold and 0.5-fold in cells derived from both donors 48 and 51 treated with high concentrations of resveratrol (48: p < 0,01, 51: p < 0,01). In contrast *P16* expression levels became significantly downregulated/ 0.62-fold in UdRPCs derived from the 48-year-old donor with the low concentration and significantly upregulated 8.8-fold when cells were treated with the high concentrations of resveratrol (48: p < 0,01). Next, we assessed the effects of resveratrol on the stem cell characteristics by measuring the expression level of the renal progenitor marker SIX2 and CD133 (fig. 4a+f+supplementary fig. 3b) as well as the occurrence of the aberrant Lamin A splicing variant progerin (fig. 4e+i). In both cell cultures we found a non-significant upregulation/ 0.35-fold and 0.16-fold of *SIX2* mRNA (48: p < 0,31 and 51: p < 0,27) when cells were treated with the low concentrations of resveratrol, while high concentrations of resveratrol significantly downregulated/ 0.99-fold and 0.45-fold (48: p < 0,01 and 51: p < 0,01) *SIX2* mRNA expression levels. In contrast for CD133 a significant upregulation/ 5.3-fold was only found in the UdRPCs derived from the 48-year-old individual (p = 0,02) when treated with the low and a significant downregulation/ 0.83-fold (p < 0,01) when treated with the high concentrations of resveratrol. UdRPCs derived from the 51-year-old individual showed no difference in CD133 mRNA expression when treated with the low and a slightly but not significant (p = 0,2) downregulation/ 0.31-fold of CD133 when treated with the high concentration of resveratrol. The aberrant Lamin A splicing variant progerin could not be detected in the UdRPCs derived from the 48-year-old individual treated with 0 μM, 0,1 μM or 1 μM of resveratrol, but alternate splicing of Lamin A occurred when cells were treated with the 2,5 μM resveratrol solution. In contrast, in the UdRPCs derived from the 51-year-old individual the two aberrant splicing variants were detected in all samples, but in cells treated with low concentrations of resveratrol the intensity of one of the detected bands became much weaker. Next, we performed Western blot analysis of the downstream targets of SIRT1, namely AKT and GSK3b (fig. 4d+h+ supplementary fig. 3a). We found that in the low DNA damage UdRPCs derived from the 48-year-old individual GSK3b-phosphorylation (p = 0,04) was significantly increased by 50% when the cells were treated with the 1μM solution of resveratrol. Furthermore, within cells from the 48-year-old individual treated with the 2.5μM solution of resveratrol GSK3b phosphorylation was found to be significantly decreased by 100% (p < 0,01). In contrast AKT phosphorylation was found to be significantly decreased when cells were treated with either of the resveratrol solutions by 80% and 90% respectively (p < 0,01). In the high DNA damage UdRPCs derived from the 51-year-old individual no significant changes in AKT or GSK3b phosphorylation were observed. Finally, we evaluated the effects of the resveratrol treatment on the proliferative capacity of UdRPCs (fig.4c+g). After 24h cells from both donors treated with the 2.5 μM solution of resveratrol showed a significant decrease in cell number (p < 0,01) (fig.4g). In contrast only the UdRPCs derived from the 48-year-old individual showed a significant increase in cell number after 24h of 0,1 and 1μM resveratrol treatment (fig.4c).

**Figure 4:**
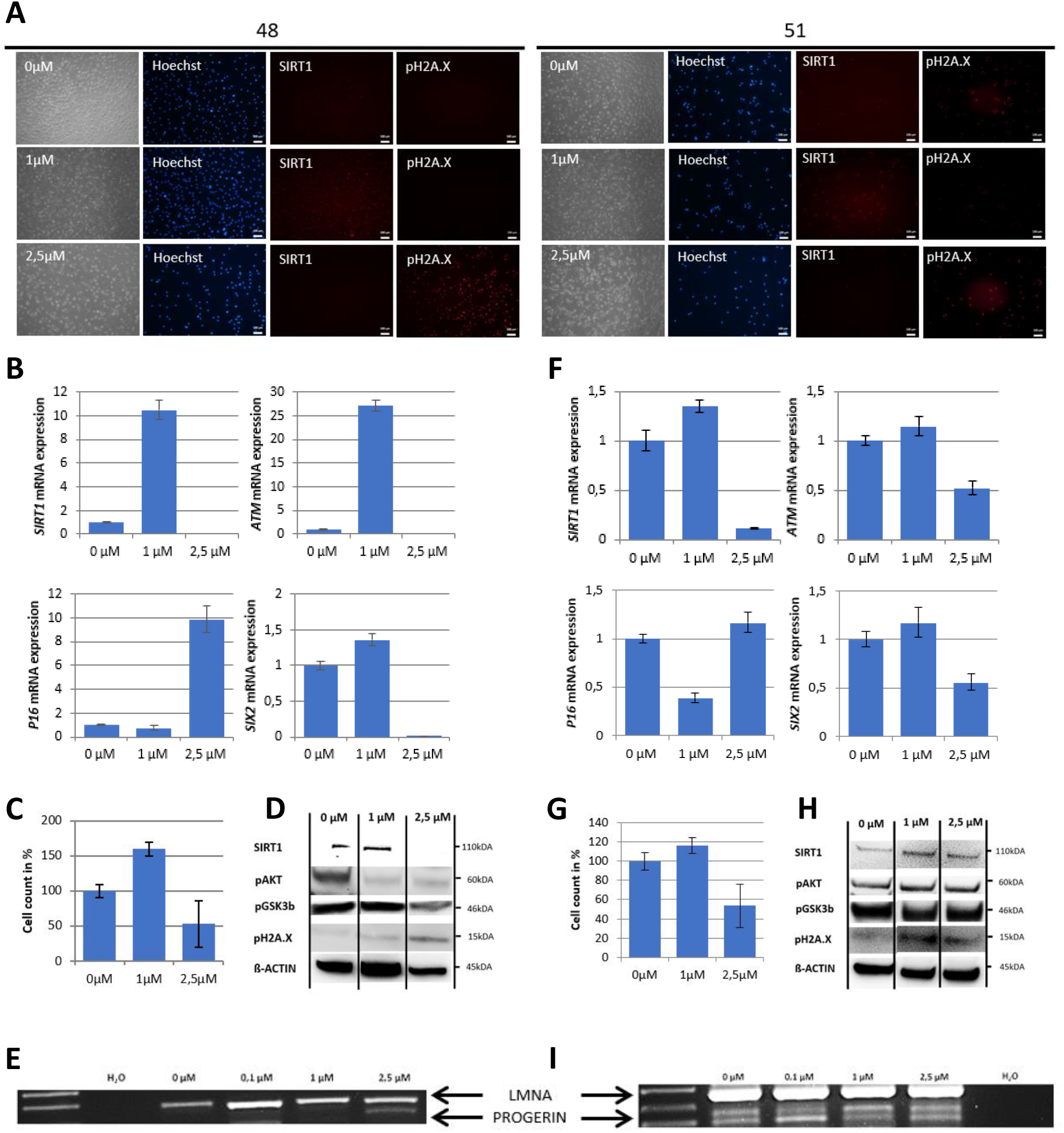
The SIX2/SIRT1/AKT/GSK3b network can be activated by resveratrol and regulates the cell fate of UdRPCs. Cell culture medium of UdRPCs was supplemented with different concentrations of resveratrol for 24h. Activation of *SIRT1* and phosphorylation of H2A.X were monitored by immunofluorescence-based detection (A) (scale bars: 100 µm). mRNA expression of *SIRT1, ATM, P16* and *SIX2* was determined by quantitative real-time PCR (B). Cell growth was evaluated after 24h of resveratrol treatment as depicted (C+G). RT-PCR analysis reveal progerin transcripts in UdRPCs treated with high concentrations of resveratrol (E+I). Relative protein expression normalized to ß-ACTIN for SIRT1 and relative protein phosphorylation for AKT, GSK3b and pH2A.X was detected by Western blot (D+H).

### DNA damage induces an aging phenotype by downregulation of SIRT1

DNA damage can trigger four potential outcomes namely, transient cell cycle arrest coupled with DNA repair, apoptosis, senescence or cell differentiation [9]. SIRT1 has been reported to participate in all of the mentioned biological processes, so we tested the effect of endogenous induced DNA damage on SIRT1 expression in UdRPCs, expressing high levels of SIRT1 [20–22]. Bleomycin has been recognized as potent inducer of DBS for many years [55, 56], so we prepared final concentrations of 1μM resveratrol, 30μg/ml Bleomycin and a combination of both substances directly in the cell culture medium and treated the UdRPCs derived from the 27-year-old individual for 24h. Interestingly while resveratrol treated UdRPCs showed a significant increase/ 0.41-fold in *SIRT1* mRNA expression (p < 0,01), upon resveratrol treatment (fig. 5b), we found SIRT1 protein expression to be unchanged between resveratrol treated and control cells (fig. 5c and supplementary fig. 4b). In contrast, cells treated with the combination of resveratrol and bleomycin showed no changes in mRNA expression, while bleomycin alone showed a significant decrease/ 0.21-fold in SIRT1 mRNA as well as protein expression (p < 0,01) (fig. 5c and supplementary fig. 4b). Immunofluorescence-based detection of H2A.X phosphorylation revealed no beneficial effect of the reseveratrol treatment alone, with 34% (control) and 33% (1μM resveratrol) of cells being positive for DBS. In contrast UdRPCs treated with either Bleomycin alone or the combination of 1μM resveratrol and 30μg/ml Bleomycin showed a significant increase (p < 0,01), with 78% and 59% of cells being positive for pH2A.X expression (fig. 5a). Furthermore, Western blot analysis of H2A.X phosphorylation normalized either to ß-actin or SIRT1 expression revealed a significant increase (p < 0,01) of DBS in cells treated with Bleomycin (fig. 5c). In contrast H2A.X phosphorylation could not be detected in either the control, the resveratrol, and the resveratrol + Bleomycin treated samples (fig. 5c). Strikingly the increased H2A.X phosphorylation was accompanied by a significant increase/ 9.23-fold of *P16* mRNA expression, while for all other conditions expression levels were found to be unchanged (p < 0,01). Surprisingly *ATM* mRNA expression levels were also found to be unchanged, with a slight downregulation in the resveratrol treated samples. Next, we assessed the effects of the resveratrol and/or Bleomycin on the stem cell characteristics of UdRPCs by measuring the expression level of the renal progenitor marker *SIX2* and *CD133* (fig.5 b). In accordance with our previous data, UdRPCs treated with resveratrol alone and in combination with Bleomycin showed an upregulation of *SIX2* (1.38-fold) and *CD133* (4-fold) mRNA levels. While this upregulation was found to be not significant when cells were treated with the combination of Resveratrol and Bleomycin (p = 0,25), the upregulation became highly significant by resveratrol treatment alone (p = 0,01). In contrast bleomycin treatment alone slightly downregulated mRNA expression levels of *SIX2* and *CD133* (fig. 5b and supplementary fig. 4a). Finally, we performed western blot analysis of the downstream targets of SIRT1, namely AKT and GSK3b (fig. 5c and supplementary fig. 4b). While AKT phosphorylation was found to be unchanged in resveratrol alone treated cells, the UdRPCs treated with the combination of resveratrol and bleomycin (by 74%) and the bleomycin alone (by 44%) treated cells showed a significant decrease (p < 0,01). Interestingly GSK3b phosphorylation was found to be significantly downregulated in UdRPCs derived from the 27-year-old individual under all culture conditions (by 60% with Resveratrol, by 89% with Resveratrol and Bleomycin and 93% with Bleomycin) (fig. 5b and supplementary fig. 4a).

**Figure 5:**
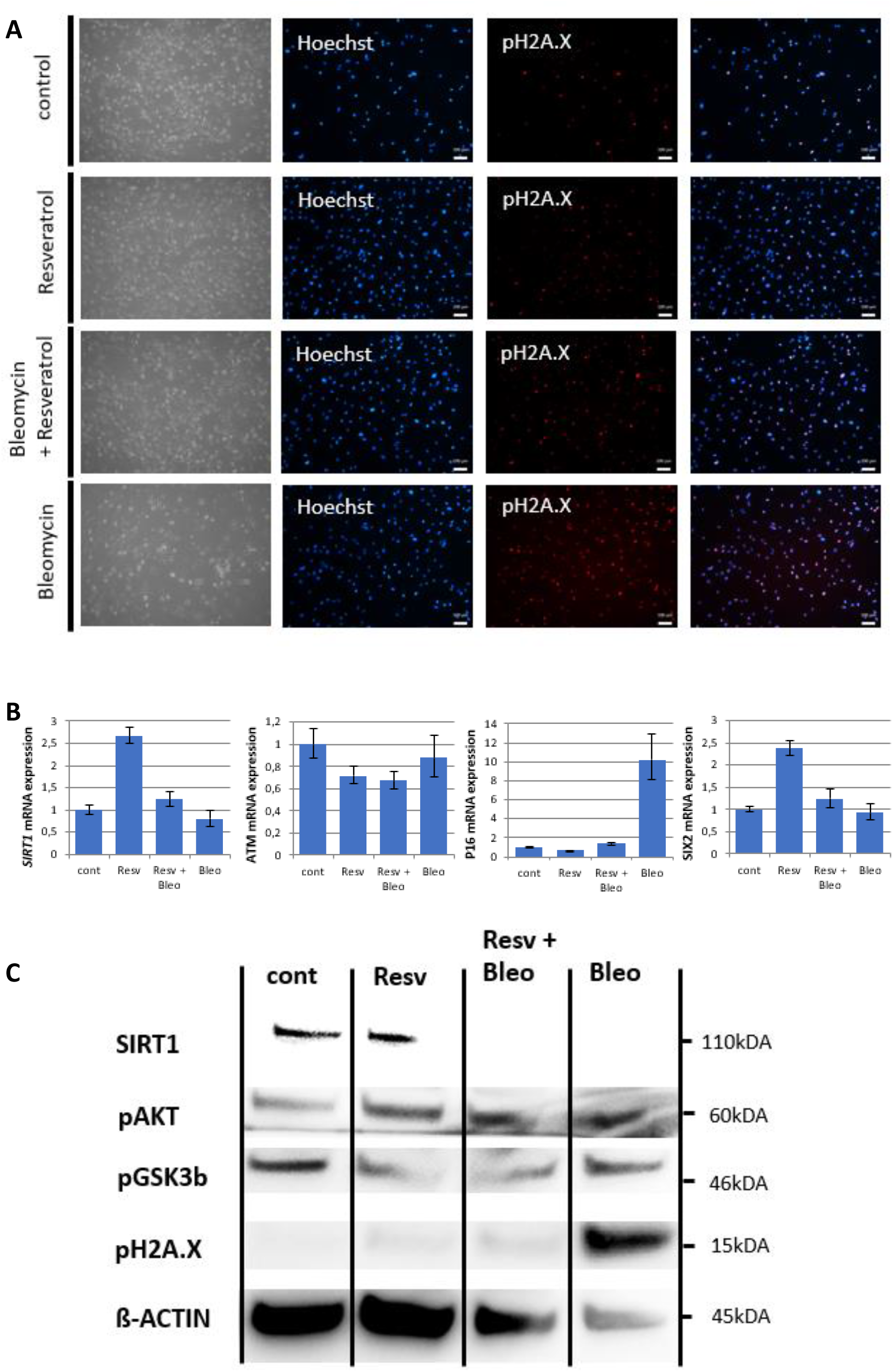
DNA damage induces an aging phenotype by downregulation of SIRT1. Cell culture medium of UdRPCs was supplemented with 1μM of resveratrol and or 30 μg/ml of Belomycin for 24h. Phosphorylation of H2A.X was monitored by immunofluorescence-based expression analysis (A) (scale bars: 100 µm). mRNA expression of *SIRT1, ATM, P16* and *SIX2* was determined by quantitative real time PCR (B). Relative protein expression normalized to ß-ACTIN for SIRT1 and relative protein phosphorylation for AKT, GSK3b and pH2A.X was detected by Western blot (C).

## Discussion

MSCs show a decline of self-renewal capacity and immunosuppressive properties with increased donor age and in vitro expansion [40, 42, 57]. In the present manuscript we provide evidence that UdRPCs directly isolated from human urine show typical hallmarks of aging when obtained from elderly donors. Our transcriptome data reveals the upregulation of genes involved in cellular senescence (*CDKN1A, CDKN2A* and *CDKN2D*) and inflammation (*CXCL1, IL6* and *IL8*) with increased donor age. In particular, the cell cycle regulator P16 (or *CDKN2A*) is believed to play a crucial role in mediating cellular senescence and preventing tumour growth [58, 59]. Furthermore, *P16* expression has been linked to the extension of normal cellular lifespan [60]. In contrast our transcriptome data revealed a negative correlation between the donor age and the expression of genes involved in genomic instability (*ATM, MSH6, TFEB* and *XRCC1*), epigenetic alterations (*SETDB1, KDM6A, EZH1, SIRT1, SETDB2, HDAC6, HDAC4, KDM4B* and *SIRT3*) and genes involved in deregulation of nutrient sensing of the one carbon-, cysteine- and methionine-metabolic pathways (*SHMT1, SHMT2* and *MAT2B*).

Somatic cells acquire mutations and other forms of DNA damage as mammals age with four potential outcomes for the affected cell namely, transient cell cycle arrest coupled with DNA repair, apoptosis, senescence or cell differentiation [9]. UdRPCs derived from aged donors show increased phosphorylation levels of Histone 2A (pH2A.X), which is an established biomarker of DNA-damage at DBS [47]. This increased amount of DBS is accompanied by a downregulation of *ATM* and lower phosphorylation levels of AKT and GSK3b. AKT signalling has been recognized to be positively affected by ATM [13] and needed for DBS repair [61]. Furthermore, a “stemness checkpoint” controlled by ATM has been suggested. Hereby, DBS initiated ATM signalling maintains MSCs and blocks differentiation [9]. This proposed “stemness checkpoint” is also reflected in our data. UdRPCs derived from young donors show low level of DNA damage accompanied with high expression levels of *ATM* and stem cell markers *CD133* and *SIX2*. Furthermore, these cells also show high phosphorylation levels of AKT and GSK3b. In contrast UdRPCs derived from elderly donors show the direct opposite expression patterns for all the mentioned factors. Of note, GSK3b inhibition has already been linked to kidney progenitor differentiation [34]. This hypothesis that UdRPCs derived from aged donors might be more prone to differentiation is strengthened by the increased amount of the aberrant splicing form of Lamin A (progerin). Since Progerin has been recognized as a marker of premature terminal differentiation and/or senescence [43].

The physiological activity of several sirtuin family members has been connected to the regulation of life span of lower organisms such as Caenorhabditis elegans [1], and Drosophila melanogaster [28] as well as mammals [29]. SIRT1 is the mammalian ortholog of the yeast silent information regulator (Sir2) and as a NAD^+^-dependent class III histone deacetylase with a wide variety of target molecules. Therefore its deacetylase activity has been linked to many biological processes connected to aging, examples-apoptosis, cell differentiation, development, stress response, metabolism, and tumorigenesis [20–22]. Interestingly, SIRT1 is recruited to DBS by ATM and is required for DNA damage repair [30]. Our transcriptome data reveals a downregulation of SIRT1 in UdRPCs derived from elderly donors. By applying genomic bisulfite sequencing we show that the SIRT1 promoter is methylation sensitive and found to be hypermethylated in UdRPCs derived from an elderly donor. It is well known that for the preservation of an unmethylated promoter DNA-methyltransferases must be excluded from the 5’-regulatory regions, which is strongly promoted by the binding of transcription factors. If a gene becomes transcriptional inactive this can lead to the progressive methylation within the 5’-regulatory region [62]. Elevated levels of genotoxic substances have been linked to increased DNA adducts, higher amounts of DNA damage and increased levels of DNMT1 expression [63]. Therefore, it is tempting to speculate that the DNA methylation changes found within the SIRT1 promoter might be a direct consequence of the increased levels of DBS in UdRPCs derived from elderly donors.

Furthermore, it has been recognized that DNA damage *in vitro* results in decreased SIRT1 activity [31] and that SIRT1 expression is reduced in senescent mesenchymal stem cells (MSCs), while its overexpression delays the onset of senescence and the loss of differentiation capacity [33]. When UdRPCs are treated with genotoxic substances (e.g., high doses of resveratrol or bleomycin), we observed a complete down-regulation of *SIRT1* mRNA and protein expression. Furthermore, high doses of genotoxic substances caused upregulation of *CDKN2A* accompanied with increased phosphorylation levels of H2A.X. Furthermore, the expression of stem cell markers *SIX2* and *CD133*, as well as the phosphorylation levels of AKT and GSK3b were found to be exclusively down-, while progerin expression was up-regulated. Strikingly, when UdRPCs are treated with low concentrations of resveratrol, which has been recognized as a potent activator of SIRT1 [52, 53], the mentioned changes within the UdRPCs could be partially reversed. Consistent in all treated UdRPCs, resveratrol caused an upregulation of *SIRT1* mRNA and protein, which was accompanied by the transcriptional upregulation of the stem cell markers *CD133* and *SIX2*, while *P16* expression was consistently downregulated. Furthermore, dependent on the accumulated DNA damage in the UdRPCs, resveratrol treatment induced an upregulation of GSK3b phosphorylation, which we conclude might enhance the self-renewal and proliferation capacity of the treated cells. This causative correlation between increased SIRT1 expression and cellular differentiation has been shown in mesenchymal stem cell models during neuronal differentiation [64]. Additionally, increased SIRT1 protein expression was found to be protective against DBS, even when cells were treated with bleomycin. Our results highlight the importance of SIRT1 in DNA damage repair recognition in UdRPCs and ultimately the control of differentiation by regulating the activation of GSK3b. Furthermore, UdRPCs can be distinguished into SIRT1 high and low expressing UdRPCs, rendering the cells with low expression levels more vulnerable to endogenous noxae. This might accelerate the accumulation of DNA damage and ultimately the accumulation of aging associated hallmarks.

To our knowledge this is the first study that reports a physical interaction of the renal progenitor marker SIX2 with the SIRT1 promoter region. The transcription factor SIX2 is needed to maintain renal progenitor cells during kidney organogenesis [34]. In UdRPCs derived from elderly donors we found a decrease in *SIX2* as well as *SIRT1* mRNA, while cells derived from young donors showed high expression levels accompanied with the already discussed consequences for cellular differentiation. This makes it tempting to speculate that both SIX2 and SIRT1 are needed to maintain self-renewal in UdRPCs and that both factors positively regulate each other. This is further strengthened by the fact that both other factors that we used as positive controls for our pull down experiment have been reported to either maintain self-renewal of nephron progenitor cells (OSR1 [50]) or to participate in the developmental process of kidney organogenesis (GDNF [51]). Upon genotoxic stress either SIX2 or SIRT1 might be aberrantly regulated with the consequence of even elevated levels of DBS leading to cellular senescence or differentiation and ultimately carcinogenic transformation. This mode of action has been proposed for the etiology of bladder cancer formation in the PrimeEpiHit hypothesis [37]. Of note, our transcriptome data revealed a downregulation of genes associated with the carbon-, cysteine- and methionine-metabolic pathways, including *MAT2B*. The promoter region of this gene has been identified to contain two resveratrol binding pockets and gets activated by resveratrol in a time- and dose-dependent manner [54]. A comparison with the cancer genome atlas TCGA reveals a downregulation of *SIRT1* and *MAT2B* for several urogenital cancer entities like, Bladder Urothelial Carcinoma, Kidney renal papillary cell carcinoma, Kidney Chromophobe, Pan-kidney cohort (KICH+KIRC+KIRP) (supplementary fig. 5).

In summary we provide evidence for a direct interaction between the renal progenitor transcription factor SIX2 and the NAD^+^-dependent class III histone deacetylase SIRT1. Both factors are needed to maintain self-renewal of CD133-positive UdRPCs. Hereby, SIRT1 is involved in deacetylation and thereby activation of protein kinase B (AKT) [23] as well as deacetylation and thereby inactivation of ß-Catenin [25]. Furthermore, AKT needs to be activated by ATM [65], even though the phosphorylation is indirect [14]. Activated AKT dephosphorylates and thereby inactivates GSK3b [65]. GSK3b phosphorylates ß-Catenin, which then become disassembled by the proteasome. Unphosphorylated and acetylated ß-Catenin is transferred to the nucleus, where it binds to TCF4 and induces nephrogenesis via activation of WNT signaling [66, 67] (fig. 6).

**Figure 6:**
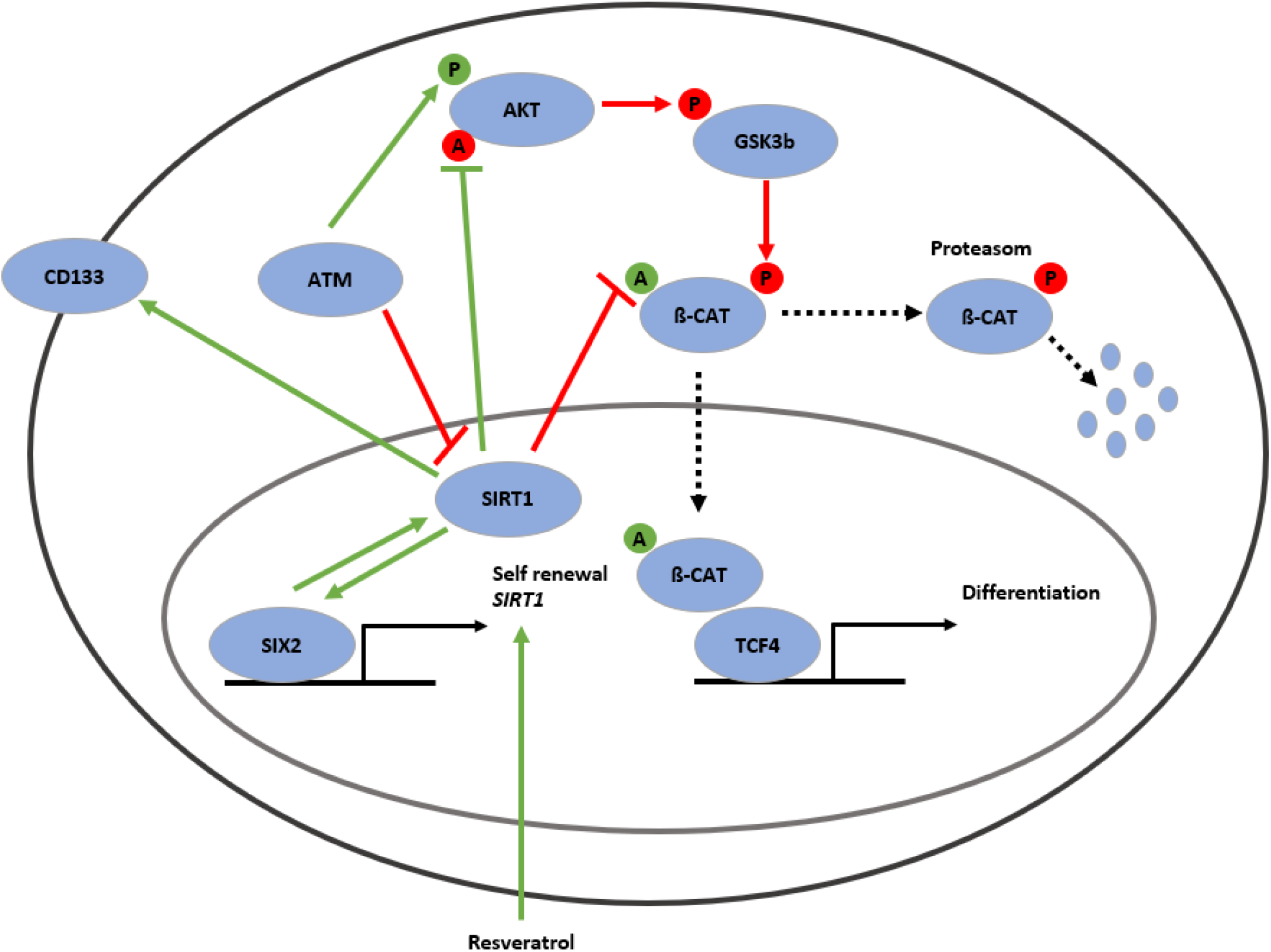
Gene regulatory network associated with the aging process in UdRPCs. SIRT1 protein is in the cell nucleus (purple circle) and can be induced by resveratrol as well as the renal progenitor-regulating transcription factor-SIX2. SIRT1 protein is positively correlated with mRNA expression of the renal stem cell markers CD133 and SIX2 and has major implications in the self-renewal of UdRPCs. SIRT1 is involved in deacetylation and thereby activation of protein kinase B (AKT) as well as deacetylation and thereby inactivation of ß-Catenin. AKT can be activated through phosphorylation by ATM. Activated AKT dephosphorylates and thereby inactivates GSK3b. GSK3b phosphorylates ß-Catenin, which then gets disassembled by the proteasome. Unphosphorylated and acetylated ß-Catenin is transferred to the nucleus, where it binds to TCF4 and induces nephrogenesis via activated WNT signaling.

## Supporting information

Table 3: differentially expressed genes udrpcs derived from differently ages donors

## Acknowledgements

Dr. Wasco Wruck for bioinformatics analysis, Chantelle Thimm, and Lisa Nguyen for technical assistance.

JA acknowledges financial support from the Medical faculty of Heinrich Heine University-Duesseldorf.

## Supporting information

Supporting information in the supplementary section or from the author.

## Conflict of interest

The authors declare no conflict of interest.

## Notes

### Competing Interest Statement

The authors have declared no competing interest.

