## supplementary information for "Crosstalk between age accumulated DNA-damage and the SIRT1-AKT-GSK3ß axis in urine derived renal progenitor cells"

Supplemental Material Table of Contents

Table 1: PCR Primers

Table 2: antibodies

Table 3: differentially expressed genes udrpcs derived from differently ages donors

Supplementary figure 1: UdRPCs show an increase of DNA damage with increase donor age

Supplementary figure 2: SIX2/SIRT1/AKT/GSK3b network is altered in UdRPCs derived from aged donors

Supplementary figure 3: The SIX2/SIRT1/AKT/GSK3b network can be activated by resveratrol and regulates the cell fate of UdRPCs

Supplementary figure 4: DNA damage induces an aging phenotype by downregulation of SIRT1

Supplementary figure 5: TCGA SIRT1 and MAT2B expression plots

Table 1: RT-qPCR Primers

| Primer name | Sequence | Annealing temperature (°C) | Product length (bp) |
|---|---|---|---|
| ATM s | 5’-AGCTCGGATGCTTTCCTCAA-3’ | 60 | |
| ATM as | 5’-CTCCATCGAGAAGGTCCACG-3’ | | |
| CD133 s | 5’-GACTTGCGAACTCTCTTGAATGA-3’ | 60 | 222 |
| CD133 as | 5’-GGTAGTGTTGTACTGGGCCAAT-3’ | | |
| CDKN2A s | 5’-CAACGCACCGAATAGTTACG-3’ | 60 | |
| CDKN2A as | 5’-AGCACCACCAGCGTGTC-3’ | | |
| GDNF s | 5’-GTCAAGAGAGGGTTTTCGGGT-3’ | 60 | 105 |
| GDNF as | 5’-ATCTTAAAGTCCCGTCCGGC-3’ | | |
| LMNA (Exon 9-12) s | 5’-GGCTGCGGGAACAGC-3’ | 60 | 500 |
| LMNA (Exon 9-12) as | 5’-CTGGCAGGTCCC-3’ | | |
| MAT2b s | 5‘-ACAGAGAGGAAGACATACCAG-3‘ | 60 | 324 |
| MAT2b as | 5‘-GTTCATTGCCAGACCAGTG-3‘ | | |
| OSR1 s | 5‘-GCTAAAGCCCCAGAGACGTG-3‘ | 60 | 80 |
| OSR1 as | 5‘-TTCGGTAGTTGCAGTGGCTT-3‘ | | |
| RPL0 s | 5‘-TCGACAATGGCAGCATCTAC-3‘ | 60 | 195 |
| RPL0 as | 5‘-ATCCGTCTCCACAGACAAGG-3‘ | | |
| SIRT1 (Exon 1-2) s | 5‘-AGGGCGAGGAGGAGGAAGAG-3‘ | 60 | 122 |
| SIRT1 (Exon 1-2) as | 5‘-GGCTCTATCCTCCTCATCACTTTC-3‘ | | |
| SIRT1 (Exon 7-8) s | 5‘-GCAGATTAGTAGGCGGCTTG-3‘ | 60 | 152 |

| | | | |
|---|---|---|---|
| SIRT1 (Exon 7-8) as | 5‘-TCTGGCATGTCCCACTATCA-3‘ | | |
| SIRT1 bseq s | 5‘-GAGGGAGGAGGGTTAGAGAG-3‘ | 55 | 341 |
| SIRT1 bseq as | 5‘-CATTATCTCCTTCCCCAACC-3‘ | | |
| SIX2 s | 5′-GGTATTATGTTTATGTTGTTTAT-3′ | 60 | 232 |
| SIX2 as | 5′-AACTAATAACTCTCCAAAATCT-3′ | | |

Table 2: antibodies

| Antigen | Company | Dilution (IF/WB) |
|---|---|---|
| Mouse ß-Actin | Cell Signaling #3700 | n.a./1:5000 |
| Rabbit pAKT (Ser473) | Cell Signaling #9271 | n.a./1:1000 |
| Rabbit pGSK3ß (Ser9) | Cell Signaling #5558 | n.a./1:1000 |
| Rabbit pH2A.X (Ser139) | Cell Signaling #9718S | 1:200/1:1000 |
| Mouse TP53 | Merck Millipore #OP43 | n.a./1:1000 |
| Mouse SIRT1 | Abcam #ab110304 | 1:200/1:1000 |
| Mouse SIX2 | Abnova #H00010736-1101 | 1:200/n.a. |
| Anti-mouse Alexa Fluor 488 | Invitrogen #A11070 | 1:500 |
| Anti-rabbit Alexa Fluor 555 | Invitrogen #A21424 | 1:500 |
| Anti-mouse HRP-labeled | Thermo Fisher Scientific #NA931 | 1:4000 |
| Anti-rabbit HRP-labeled | Cell Signaling #7074S | 1:1000 |

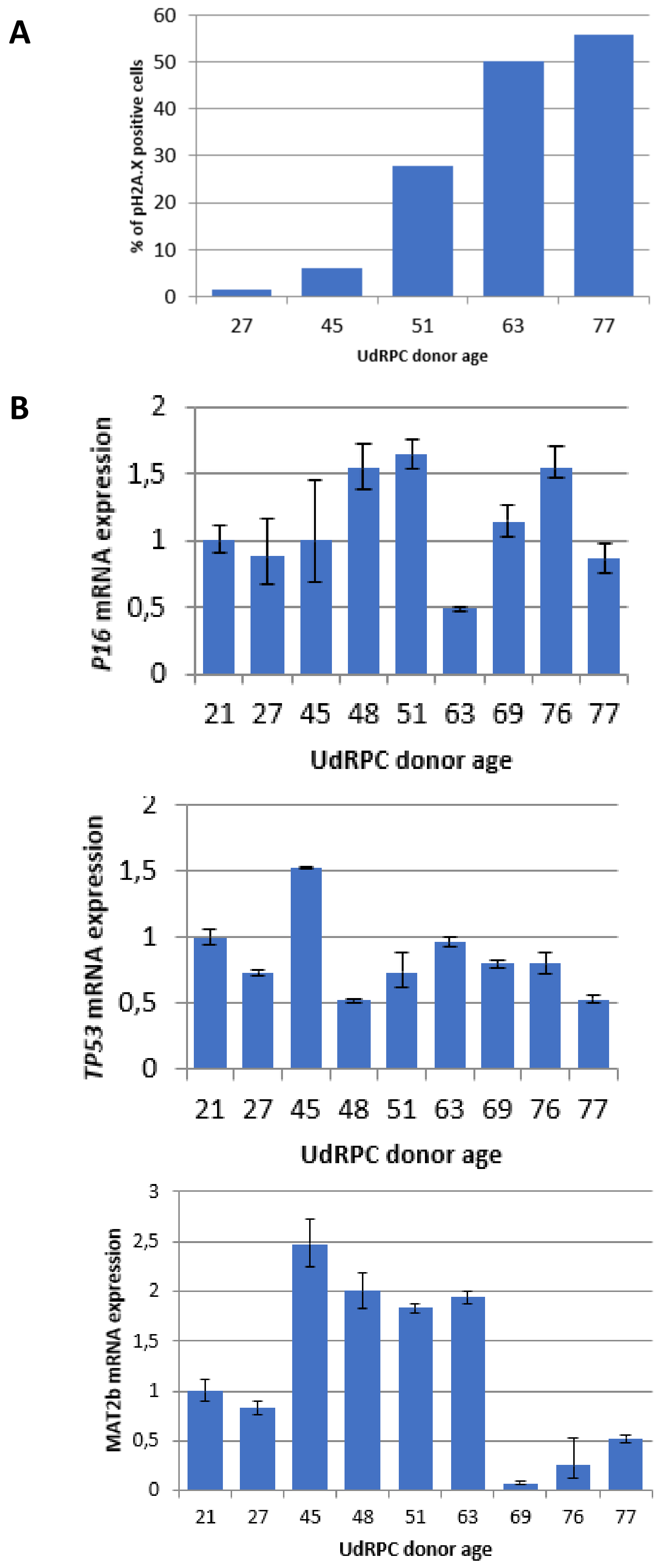


**Supplementary figure 1: UdRPCs show an increase of DNA damage with increase donor age**

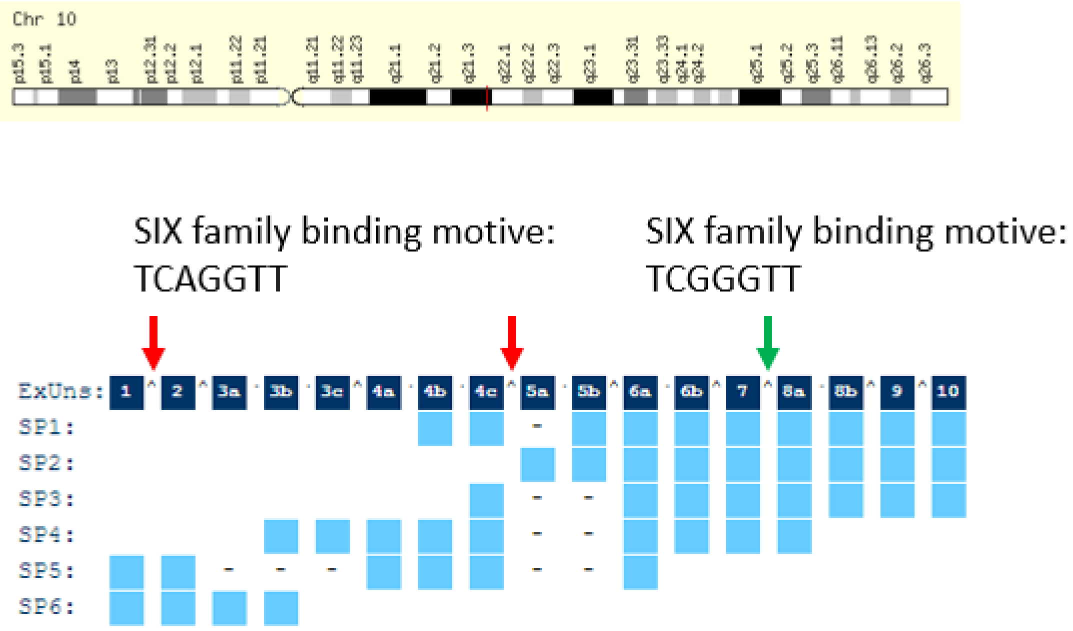


**Supplementary figure 2: SIX2/SIRT1/AKT/GSK3b network is altered in UdRPCs derived from aged donors**

**A**

**48**

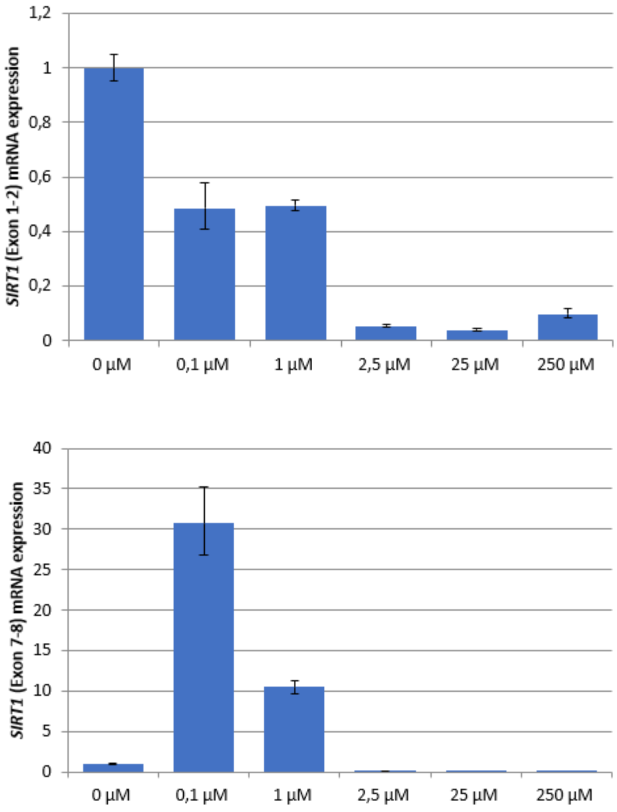

**51**

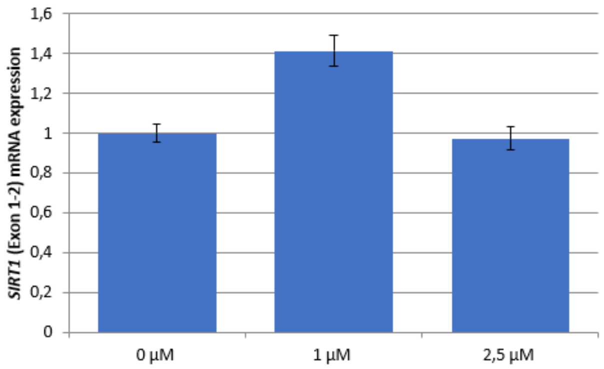
SIRT1 (Exon 1-2) mRNA expression
1,6
1,4
1,2
1
0,8
0,6
0,4
0,2
0
0 µM
1 µM
2,5 µM

48

51

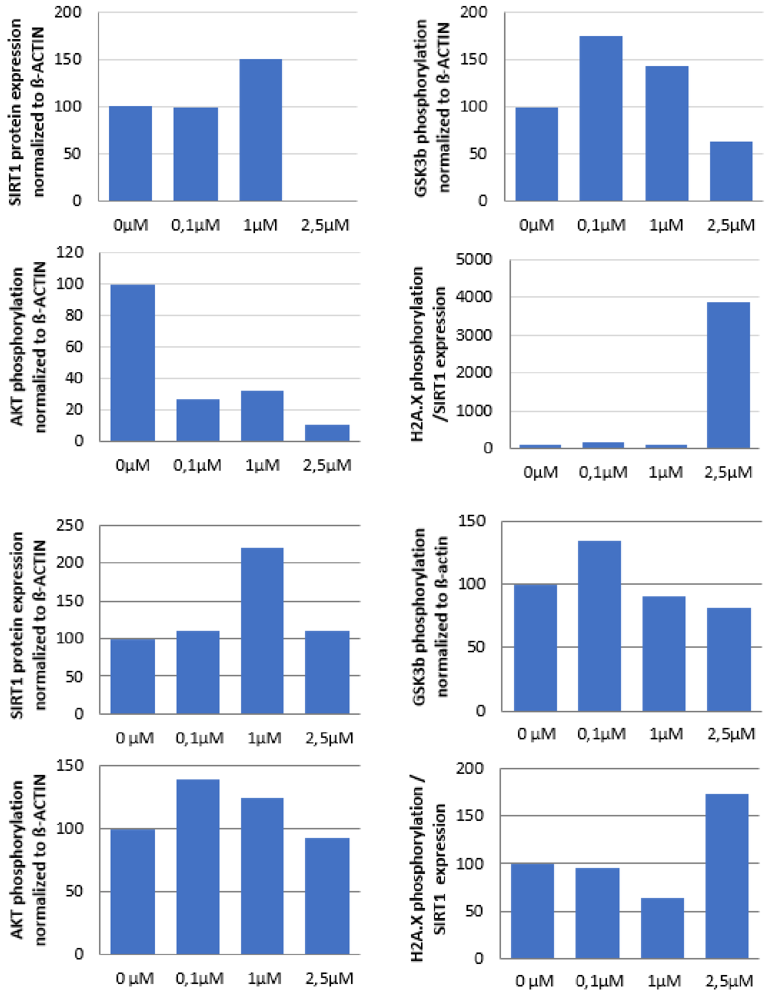
SIRT1 protein expression normalized to ß-ACTIN
GSK3b phosphorylation normalized to ß-ACTIN
AKT phosphorylation normalized to ß-ACTIN
H2A.X phosphorylation /SIRT1 expression
SIRT1 protein expression normalized to ß-ACTIN
GSK3b phosphorylation normalized to ß-actin
AKT phosphorylation normalized to ß-ACTIN
H2A.X phosphorylation / SIRT1 expression
0µM
0,1µM
1µM
2,5µM

**B**

**48**

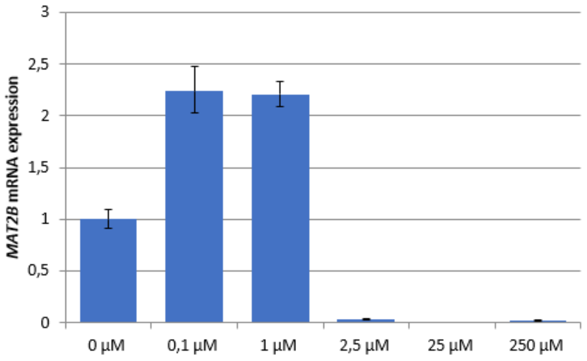
*MAT2B* mRNA expression
3
2,5
2
1,5
1
0,5
0
0 µM
0,1 µM
1 µM
2,5 µM
25 µM
250 µM

**51**

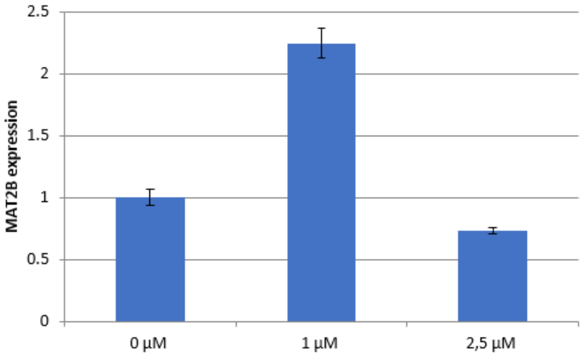
MAT2B expression
2.5
2
1.5
1
0.5
0
0 µM
1 µM
2,5 µM

**51**

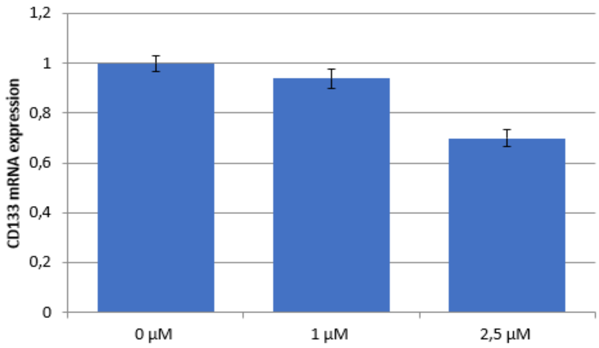
CD133 mRNA expression
1,2
1
0,8
0,6
0,4
0,2
0
0 µM
1 µM
2,5 µM

**48**

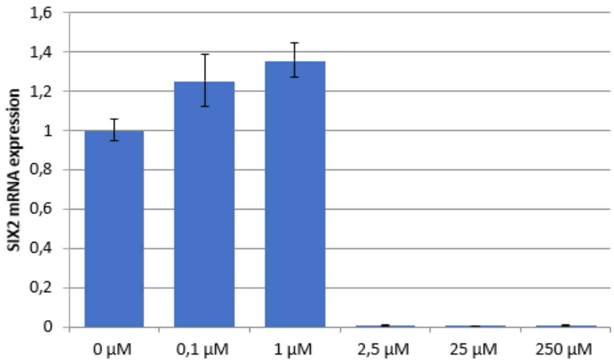
SIX2 mRNA expression
1,6
1,4
1,2
1
0,8
0,6
0,4
0,2
0
0 µM
0,1 µM
1 µM
2,5 µM
25 µM
250 µM

48

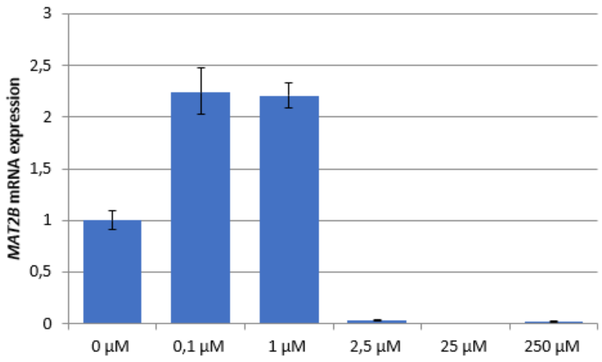


48

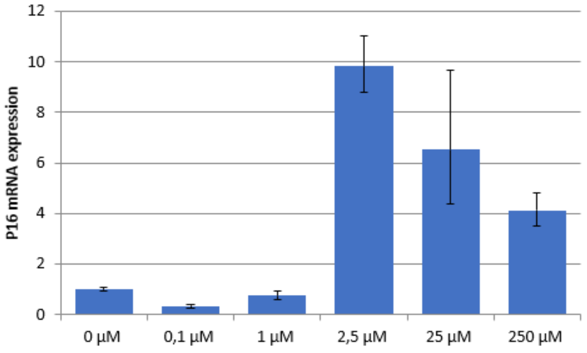


48

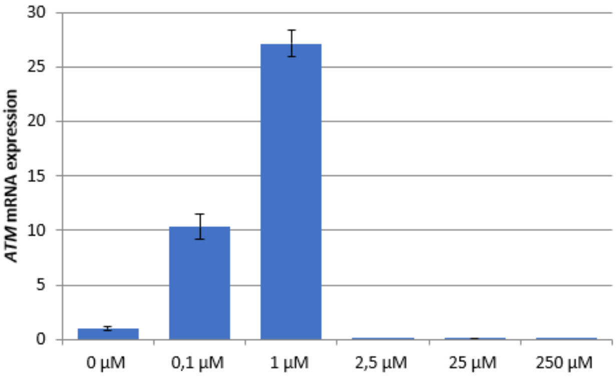


48

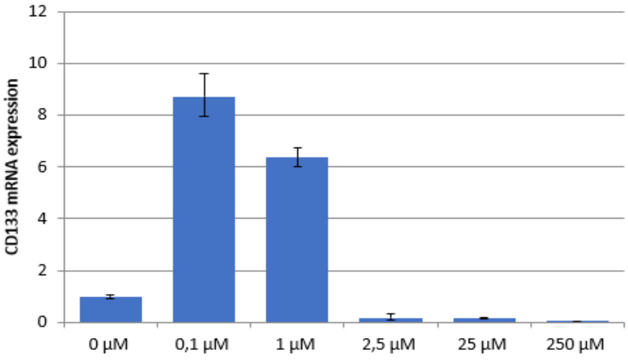


**Supplementary figure 3: The SIX2/SIRT1/AKT/GSK3b network can be activated by resveratrol and regulates the cell fate of UdRPCs**

**A**

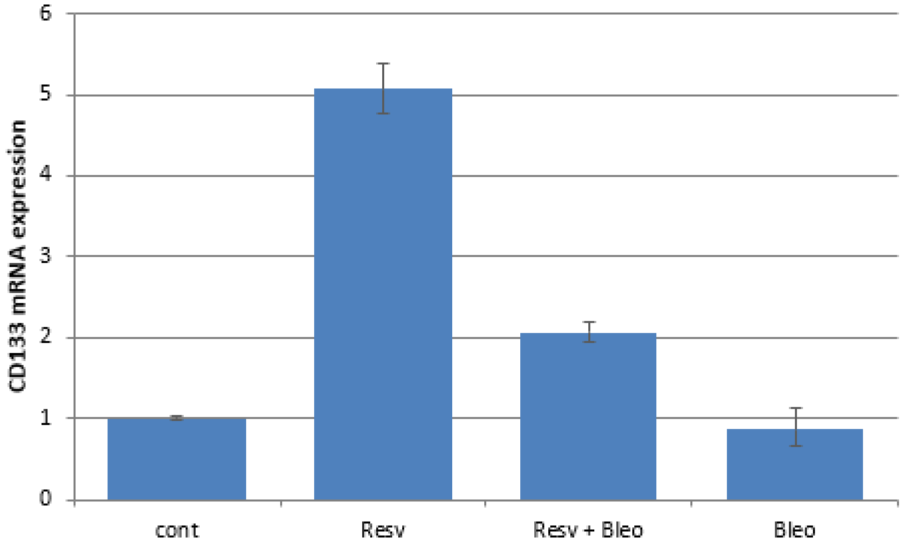


**B**

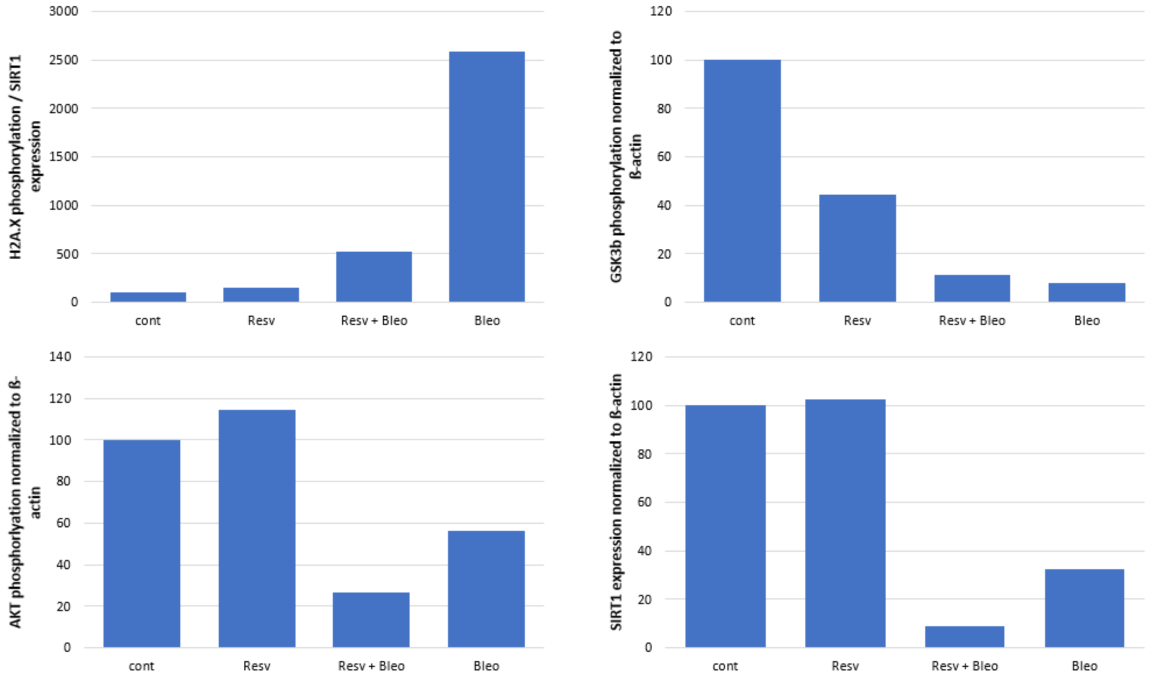


**Supplementary figure 4: DNA damage induces an aging phenotype by downregulation of SIRT1**

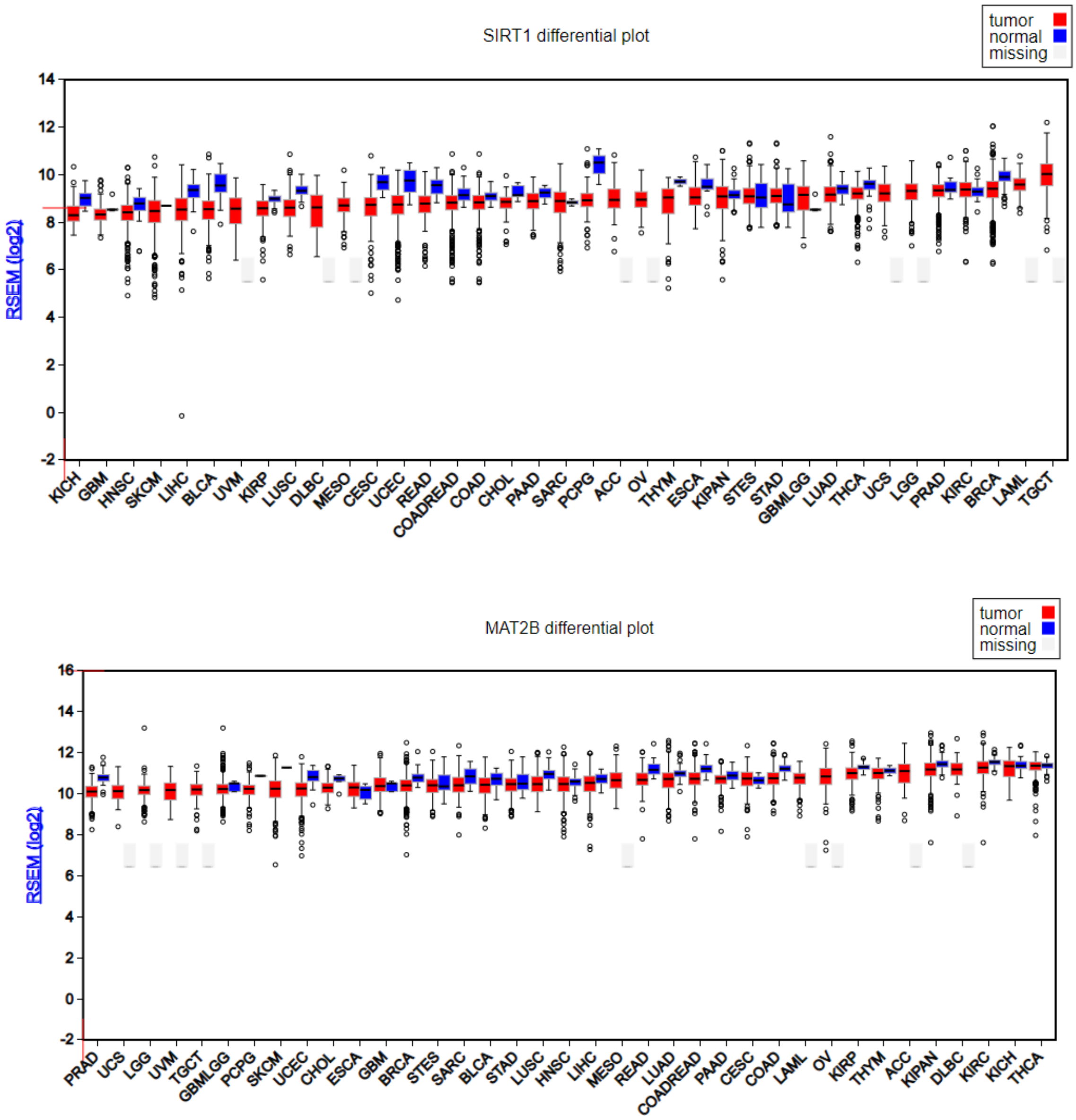


**Supplementary figure 5: TCGA SIRT1 and MAT2B expression plots.**
